# Coordinated Changes in Gene Expression Kinetics Underlie both Mouse and Human Erythroid Maturation

**DOI:** 10.1101/2020.12.21.423773

**Authors:** Melania Barile, Ivan Imaz-Rosshandler, Isabella Inzani, Shila Ghazanfar, Jennifer Nichols, John C. Marioni, Carolina Guibentif, Berthold Göttgens

**Affiliations:** Department of Haematology, University of Cambridge, CB2 0AW Cambridge, UK; Wellcome-Medical Research Council Cambridge Stem Cell Institute, University of Cambridge, CB2 0AW Cambridge, UK; University of Cambridge Metabolic Research Laboratories and MRC Metabolic Diseases Unit, CB2 0QQ Cambridge, UK; Cancer Research UK Cambridge Institute, University of Cambridge, CB2 0RE Cambridge, UK; Department of Physiology, Development and Neuroscience, University of Cambridge, CB2 3DY Cambridge, UK; Wellcome Sanger Institute, Wellcome Genome Campus, CB10 1SA Cambridge, UK; European Molecular Biology Laboratory, European Bioinformatics Institute (EMBL-EBI), Wellcome Genome Campus, CB10 1SD Cambridge, UK; Sahlgrenska Center for Cancer Research, Department of Microbiology and Immunology, University of Gothenburg, 413 90 Gothenburg, Sweden

**Keywords:** RNA velocity, gastrulation, erythropoiesis, Gata1

## Abstract

**Background:** Single cell technologies are transforming biomedical research, including the recent demonstration that unspliced pre-mRNA present in single cell RNA-Seq permits prediction of future expression states. Here we applied this ‘RNA velocity concept’ to an extended timecourse dataset covering mouse gastrulation and early organogenesis.

**Results:** Intriguingly, RNA velocity correctly identified epiblast cells as the starting point, but several trajectory predictions at later stages were inconsistent with both real time ordering and existing knowledge. The most striking discrepancy concerned red blood cell maturation, with velocity-inferred trajectories opposing the true differentiation path. Investigating the underlying causes revealed a group of genes with a coordinated step-change in transcription, thus violating the assumptions behind current velocity analysis suites, which do not accommodate time-dependent changes in expression dynamics. Using scRNA-Seq analysis of chimeric mouse embryos lacking the major erythroid regulator *Gata1*, we show that genes with the step-changes in expression dynamics during erythroid differentiation fail to be up-regulated in the mutant cells, thus underscoring the coordination of modulating transcription rate along a differentiation trajectory. In addition to the expected block in erythroid maturation, the *Gata1*^-^ chimera dataset revealed induction of PU.1 and expansion of megakaryocyte progenitors. Finally, we show that erythropoiesis in human fetal liver is similarly characterized by a coordinated step-change in gene expression.

**Conclusions:** By identifying a limitation of the current velocity framework coupled with *in vivo* analysis of mutant cells, we reveal a coordinated step-change in gene expression kinetics during erythropoiesis, with likely implications for many other differentiation processes.

## Background

Cellular differentiation into diverse cell types underpins all metazoan development. Moreover, cellular differentiation processes are also crucial for stem cell-mediated tissue maintenance, and their perturbation has been implicated in ageing-associated regenerative failure as well as malignant transformation (1, 2). Since cellular differentiation decisions are made at the level of individual cells, elucidation of the underlying molecular mechanisms requires the use of single cell approaches. It is no surprise therefore that recent innovations in single cell molecular profiling technologies have been embraced rapidly by developmental and stem cell biologists, with complete single cell gene expression maps now available for developing embryos of several model organisms (3-5, reviewed in 6), as well as large-scale datasets covering adult tissue homeostasis (7–9).

Comprehensive molecular profiling necessarily entails the generation of snapshot data, because cells need to be fixed to examine their molecular content. This in turn represents a major drawback for the study of differentiation processes, which commonly occur over extended timeframes via complex trajectories underpinned by intricate decision-making processes. Much excitement was therefore generated by a recent seminal study (10), which demonstrated that unspliced pre-mRNA present in scRNA-Seq datasets can be exploited to predict likely future expression states. This so-called RNA velocity concept is based on the notion that the ratio between unspliced and spliced RNA differs depending on whether a gene is in the process of being up- or downregulated. During upregulation, there is a relative increase in newly transcribed unspliced RNA, with the converse occurring during downregulation. The RNA velocity framework has rapidly gained traction across the wider single cell community, being applied across multiple experimental systems (11–13), and also extended as part of the scVelo analysis suite (14), which allows inclusion of genes whose transcript levels are not in steady state.

One system where the RNA velocity concept has particular potential is erythropoiesis, the process whereby oxygen-transporting red blood cells are generated from multipotent haematopoietic progenitors. Research into the transcriptional control processes of erythropoiesis led to several paradigmatic discoveries, including the dissection of distal transcriptional control elements (15–17), as well as antagonistic transcription factor pairings as executors of lineage choice in multipotent progenitors (18). During embryogenesis, a first so-called primitive wave of erythropoiesis occurs in the yolk sac, followed by a second definitive wave, initiated also in the yolk sac, then predominantly in the fetal liver and later in the adult bone marrow (19). The zinc finger protein Gata1 represents the archetypal erythroid transcription factor, and is required for the maturation of both primitive and definitive erythroid cells (20–23), as well as megakaryocyte maturation (24). However, the precise molecular processes affected by Gata1 deletion in early embryonic erythropoiesis have remained obscure, principally because conventional biochemical methods are unsuitable for the very small number of cells present at these early developmental stages.

Here, we have applied RNA velocity to a recently published scRNA-Seq dataset of nine sequential timepoints, spaced 6 hours apart, which encompass mouse gastrulation and early organogenesis (25). We observed that some of the inferred trajectories are incompatible with the existing biological knowledge, as well as with the real time ordering derived from the sequential sampling timepoints. For erythroid differentiation in particular, we show that failure of the velocity framework is due to a concerted increase in transcription rate of a subset of erythroid genes, midway through the red blood cell maturation trajectory. Analysis of *Gata1*^-^ chimeric embryos underscores the concerted nature of this expression boost, consistent with the notion that such concerted upregulation events may be a feature of stabilizing a given differentiated cellular state.

## Results

### Limitations of RNA velocity trajectory inference at organismal scale

To evaluate RNA velocity-based trajectory inference with a complex dataset, we applied the scVelo analysis pipeline (14) to a recently reported timecourse scRNA-Seq dataset covering mouse gastrulation and early organogenesis. This mouse gastrulation atlas contains approximately 120,000 single cell transcriptomes across nine sequential timepoints covering 37 major cell types (25). Prior to scVelo analysis, we removed extraembryonic ectoderm and extraembryonic endoderm cells, as they derive from early lineage branching events that are not covered in this dataset. We first applied scVelo to the normalised and batch corrected count matrix across all embryonic stages (Figure 1A). We observed that scVelo correctly identifies the epiblast population as the origin of the global differentiation processes that occur during gastrulation and early organogenesis. In relation to the more differentiated cell types however, there were several instances where scVelo had difficulty in capturing some of the highly complex differentiation events that occur across the entire embryo. For instance, scVelo predicted that E8.0 allantois and mesenchyme cell-types give rise to mesodermal cells from earlier timepoints rather than the E8.25/E8.5 allantoic and mesenchymal cells. Another inconsistency occurred with E8.0-E8.25 endoderm cells, which were predicted to give rise to E6.5-E7 visceral endoderm, rather than the other way round. Most noteworthy, scVelo failed to recapitulate the erythropoiesis branch, where it predicts a backwards differentiation from later to earlier populations. We next repeated this analysis using data from each individual time-point (Figure 1B; shown are E7.5 and E8.5). We saw that the pipeline accurately recapitulates known biological trajectories up to E7.5, but observed the same inconsistency from E7.75 to E8.5, with scVelo arrows pointing backwards.

**Figure 1.**
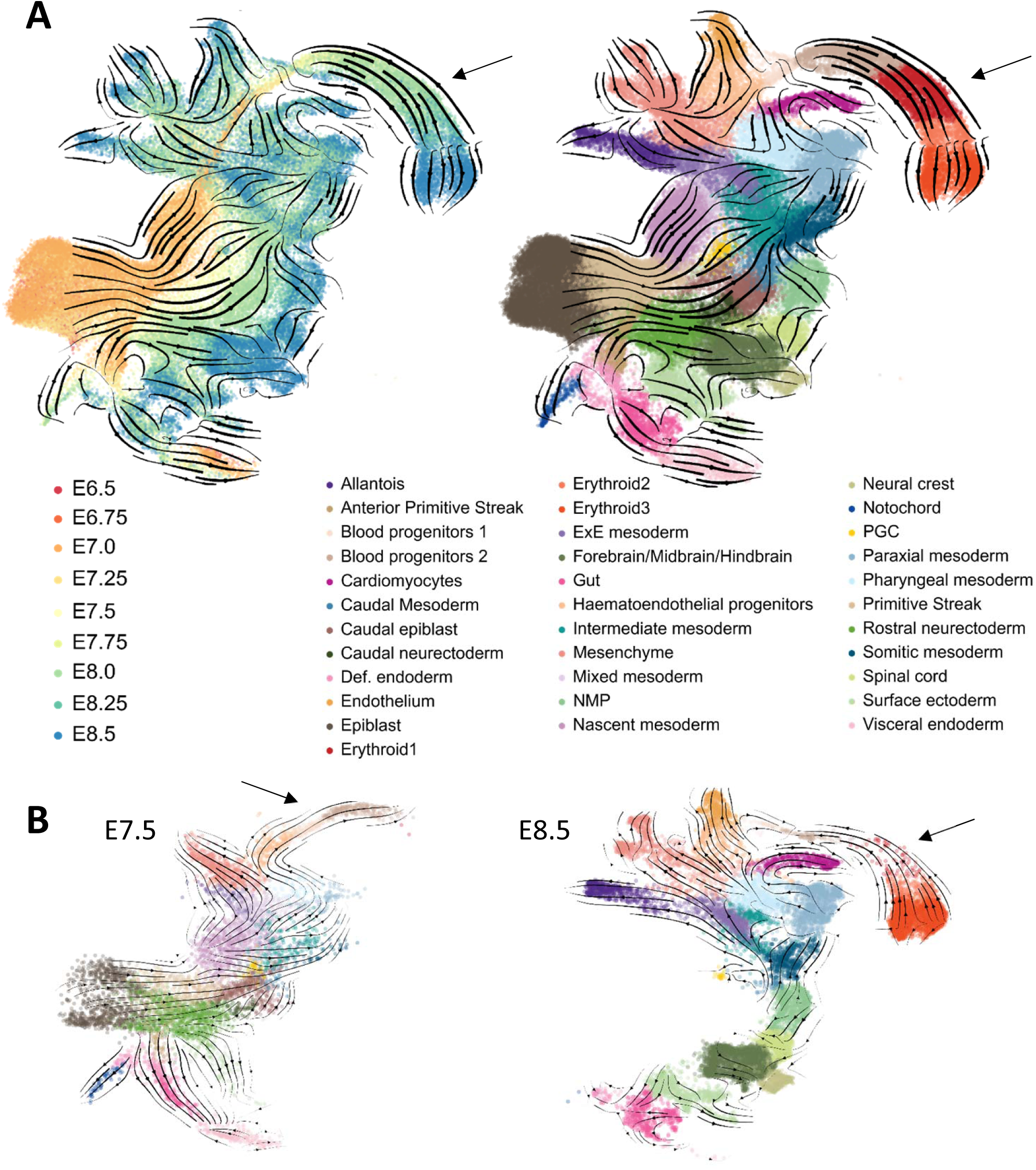
Inferring Differentiation Trajectories at organismal scale. A. Pijuan-Sala et al. (2019) layout containing single-cell transcriptomes belonging from E6.5 to E8.5, colored by sampled time-point (left) and by cell-type (right). The overlaying arrows result from applying the scVelo pipeline to the whole embryonic dataset and represent inferred developmental trajectories. Arrowheads highlight the erythroid branch, displaying scVelo trajectory predictions that are inconsistent with real-time sampling. B. Pijuan-Sala et al. (2019) layout highlighting single-cell transcriptomes belonging to E7.5 (left) and E8.5 (right) and colored by cell-type (see legend in A). The overlaying arrows result from applying the scVelo pipeline to these individual time-points and represent inferred developmental trajectories. Arrowheads highlight the erythroid branch.

Taken together therefore, we have identified that for erythroid development, the output of scVelo is inconsistent with the timecourse information gathered from the experimental design of the gastrulation atlas.

### Unspliced sequence reads help to discriminate between cell types

We next asked whether this issue is due to a general lack of biologically meaningful information captured in the unspliced reads. To this end, we exploited two variance-based dimensionality reduction methods, Principal Component Analysis (PCA) and Multi-Omics Factor Analysis (MOFA; 26), to interrogate how much inter-population variability is explained by the spliced and unspliced information layers, whether considered separately or together. Upon comparing PC1 and PC2 (or MOFA Factors 1 and 2), in addition to the expected lineage separation obtained using the spliced reads (Figure 2A, left panel), we could also observe a degree of lineage separation when using the unspliced reads alone (Figure 2A, middle panel). In addition, we saw a qualitatively improved separation of the different lineages when spliced and unspliced information is used in combination (Figure 2A, right panel; see Supplementary Figure 1 for further components/factors). Moreover, the MOFA factors account for 16% of variation in the spliced data and 4% of the of variation in unspliced data (Figure 2Bi). Interestingly, a closer look at the MOFA pre-processing and final outcome showed a minor overlap of genes that are highly variable with respect to spliced or unspliced counts (Figure 2Bii) and a different weight contributed by the two layers to the final factors (Figure 2Biii).

**Figure 2.**
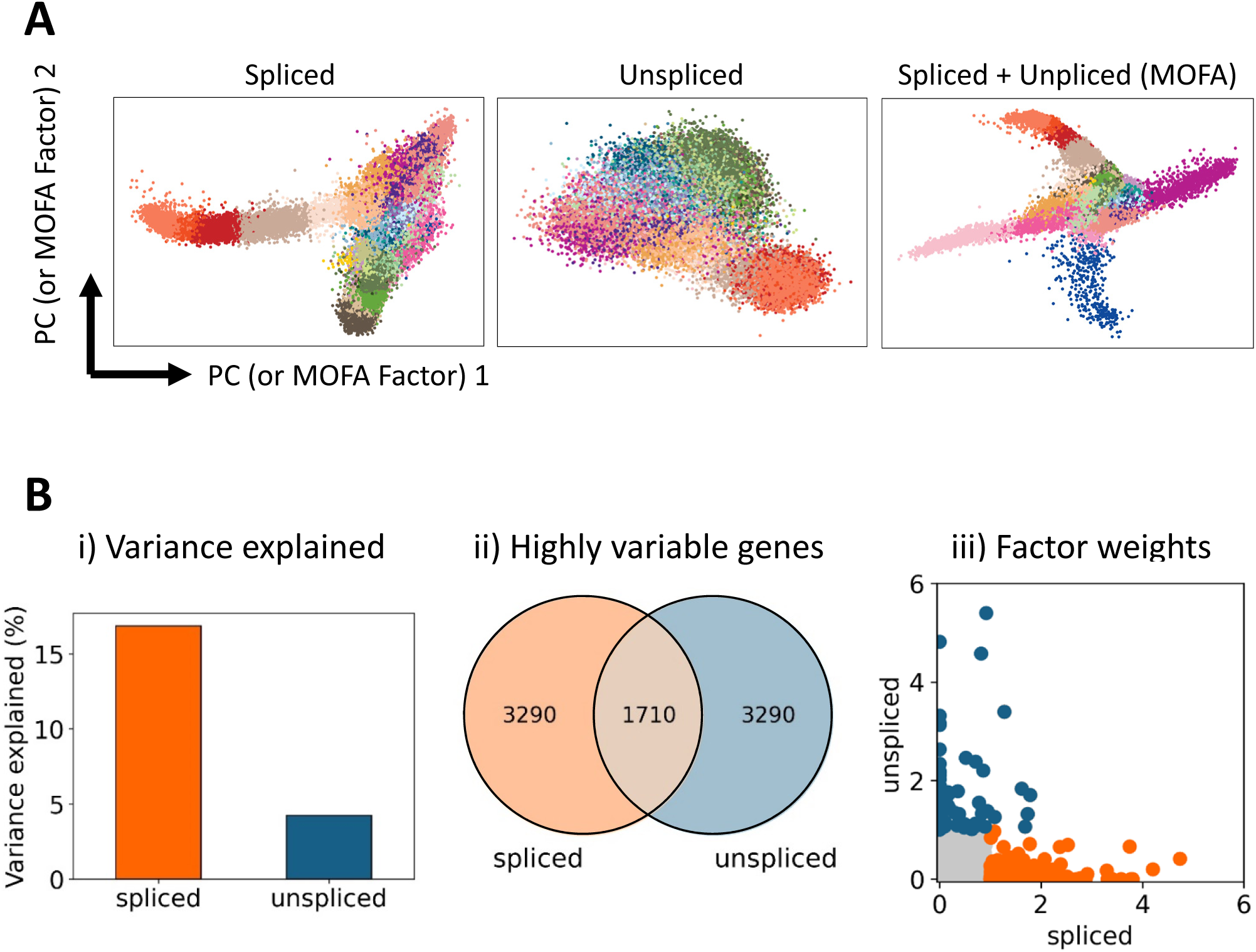
Unspliced counts contribute to explaining the variability among cell types. A. Dimensionality reduction with the first two principal components/MOFA factors using spliced reads alone (left), unspliced reads alone (middle) and both spliced and unspliced (right). Single-cell transcriptomes are colored by cell-type annotation; see Figure 1 for full legend. B. MOFA characterization of spliced and unspliced reads assessing proportion of variance explained (i), overlap in highly variable genes calculating using either spliced or unspliced reads (ii), and factor weight distributions (iii).

Multiomics factor analysis therefore not only demonstrates that the unspliced reads in the gastrulation atlas dataset contain biologically relevant information, but also suggests that integrated analysis of spliced and unspliced reads may more broadly facilitate the interpretation of complex scRNA-Seq datasets.

### Analysis of unspliced reads reveals complex expression kinetics

Having confirmed the utility of unspliced reads, we next explored whether the inability to recover real-time progression in whole embryo trajectory inference using scVelo might be related to the assumptions made by the current RNA velocity analysis tools. The derivation of gene-specific expression kinetics underpins the scVelo analysis pipeline, as illustrated by so-called phase plots that depict the amounts of spliced versus unspliced reads within a population of cells (14). If a gene is upregulated during a differentiation timecourse, cells will be placed above the diagonal between no expression and maximum expression due to the relatively larger amount of newly produced pre-mRNA during the gene induction process, while the converse is true for downregulated genes (Figure 3A). Both of these scenarios are readily captured by scVelo, with the predicted vectors of differentiation agreeing with the actual temporal progression. If a given gene however experiences an increase in transcription rate midway through a differentiation timecourse, the sudden increase in unspliced pre-mRNA will result in a phase plot that may be wrongly classified by scVelo, with predicted vectors of differentiation diametrically opposed to the true direction of differentiation (Figure 3A). This is indeed what we observed when inspecting the phase plots of the scVelo driver genes (top-likelyhood genes, Supplementary Table 1), which display a steep increase of unspliced counts in the Erythroid 3 population, leading to a reverse velocity prediction, progressing from Erythroid 3 to earlier populations (Supplementary Figure 2A).

**Figure 3.**
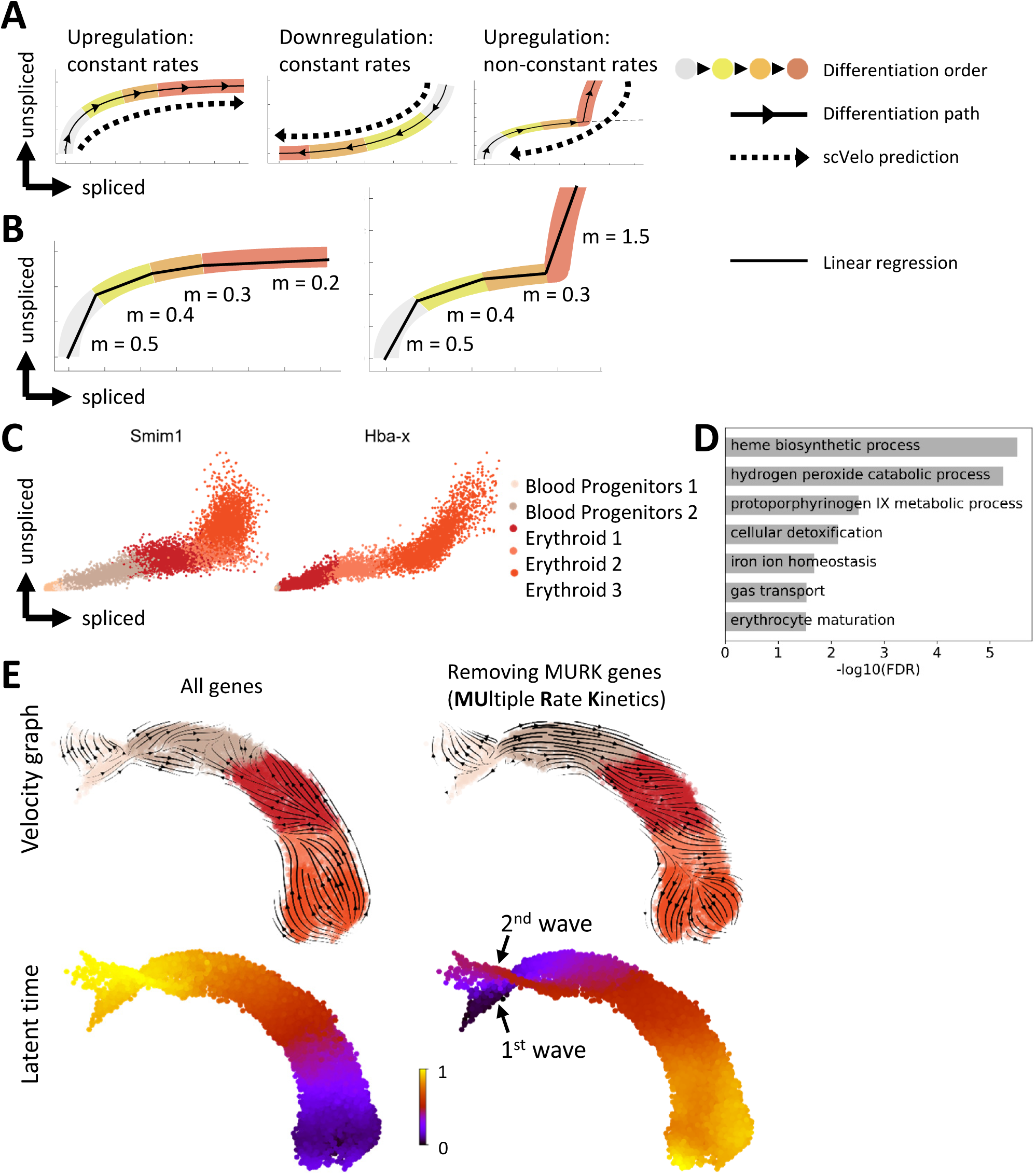
A set of genes with complex expression kinetics confounds velocity estimation in erythropoiesis. A. Illustration of phase plot representation in datasets of differentiating cell populations, and associated scVelo predictions B. Illustration of strategy for MURK gene identification C. Phase plots of representative MURK genes. X-axis: normalized imputed counts of spliced transcript; y-axis: normalized imputed counts of unspliced transcript. D. GO-term enrichment of MURK genes identified in mouse yolk sac erythropoiesis E. Zoomed-in UMAP of the erythroid branch (see Figure 1 for full UMAP) with scVelo calculations, before and after removing MURK genes identified in B. Distinct waves of embryonic erythropoiesis are visible upon MURK gene removal, highlighted with arrowheads.

We next set out to identify all genes exhibiting this rapid increase in expression levels in the Erythroid 3 population (Figure 3B). After fitting a linear regression through each population and each gene and testing whether the inferred slopes reflected the expected order based on biological knowledge, we found 89 such genes, which we termed Multiple Rate Kinetics or MURK genes. These genes included *Smim1*, coding for the Vel Blood Group Antigen (27), and *Hba-x*, where we could confirm an increase in expression kinetics using phase plots (Figure 3C).

Having identified a set of genes with a coordinated increase in expression rate midway through erythropoiesis, we next asked what function these genes might play in the broader transcriptional program of red blood cell maturation. Visual inspection of the gene list revealed it to contain archetypal red blood cell genes including the globin genes *Hba-x*, *Hbb-a1*, *Hba-a2*, *Hbb-bt*, *Hbb-bh1*, *Hbb-y* (Supplementary table 2). Unsupervised gene ontology analysis confirmed that biological functions essential for red blood cells were highly enriched, including “gas transport” and “heme biosynthetic process” (Figure 3D).

We next removed this set of MURK genes and recalculated the RNA velocity inferred trajectories. As can be seen in Figure 3E, inferred vectors of differentiation are now in good agreement with the real time progression of erythropoiesis

The scVelo suite also calculates a so-called latent time, which represents the pseudotime ordering hidden in the spliced and unspliced dynamics, and is more powerful than previously described pseudotime inferring approaches since it incorporates both the gene dynamics and the spliced and unspliced information (14). Using the full gene set, the latent time calculation for the erythroid lineage is contrary to the know progression of erythroid differentiation (Figure 3E left panels, Supplementary Figure 2B, left panels). By contrast, removing the MURK genes results in a latent time prediction that is not only consistent with the major axis of erythropoiesis, but also identifies the two sequential inputs described previously (25), namely an early wave directly from posterior mesoderm as well as a second wave coming from yolk sac hemogenic endothelium (see Figure 3E, Supplementary Figure 2B, right panels).

Taken together therefore, this analysis shows that inconsistent RNA velocity-inferred trajectories can be remedied by the removal of genes with complex expression kinetics.

### Erythroid Multiple Rate Kinetics genes are essential for red blood cell function

To corroborate upregulation of our identified MURK genes during erythropoiesis, we interrogated a previously published dataset with transcriptomic analysis of a loss of function model for the erythropoiesis master-regulator *Gata1* (28). *In vitro* differentiation of Gata1 knock-out embryonic stem cells over-expressing human *BCL2* can produce permanently self-renewing immature erythroid progenitor cell lines. One such model, G1ER, contains a tamoxifen-inducible Gata1 transgene, the activation of which triggers erythroid maturation (29, 30; Figure 4A). Microarray-based differential gene expression was performed, comparing the uninduced and induced conditions (28). 76 of our 89 MURK genes overlapped with the genes identified by this microarray-based comparison. Of those, 64 were upregulated, of which 55 showed strong upregulation, 4 were downregulated, and 8 showed no change in expression following induction of Gata1 in the G1ER system, demonstrating a highly significant overlap of our identified MURK genes with the G1ER-induced genes (p < 10^-24^; see Figure 4B).

**Figure 4.**
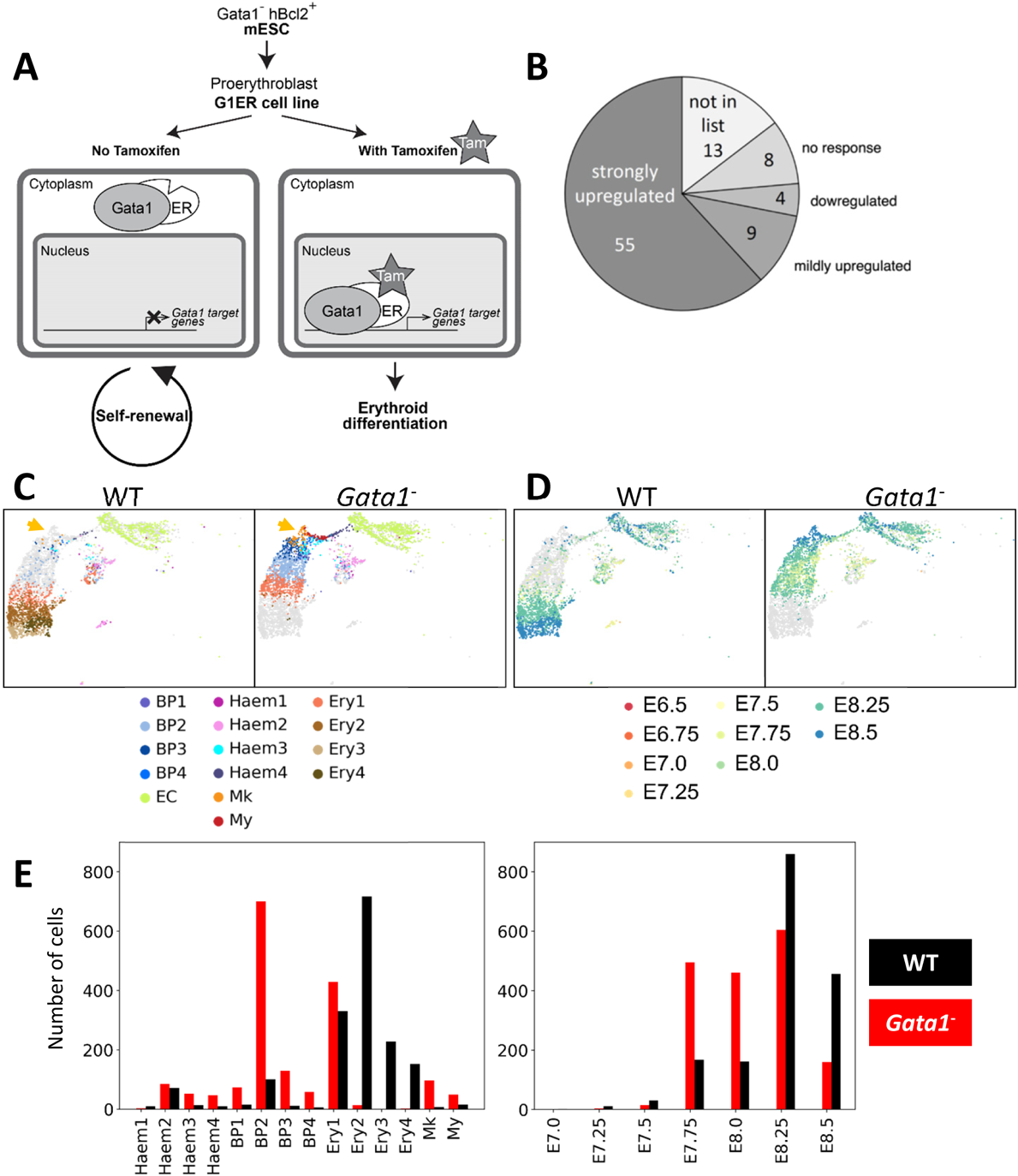
*In vivo* analysis of Gata1 function using a chimaera assay coupled with scRNA-Seq. A. Schematic of the G1ER system (29, 30) B. Behaviour of the 89 MURK genes identified in Figure 3 upon Gata1 induction in the G1ER system (28). Wu et al. report that upon Gata1 induction they obtained a total of 2769 upregulated genes, 6079 mildly upregulated, 3566 downregulated, and 3445 with no response. C. UMAPS of *Gata1*^-^ chimera cells allocated a hemato-endothelial identity colored by cell-type (sub-clusters defined in Pijuan-Sala et al. (2019) - BP: Blood Progenitors, EC: Endothelial Cells, Haem: Hemato-endothelial Progenitors, Mk: Megakaryocytes, My: Myeloid cells, Ery: Erythroid cells) and split by genotype. Orange arrowheads highlight increased population with megakaryocytic signature in Gata1^-^ fraction. D. UMAPS of *Gata1*^-^ chimera cells allocated a hemato-endothelial identity colored by sampling timepoint and split by genotype. E. Barplots with the quantification of chimera cells mapping to each hemato-endothelial lineage of the reference dataset (left) and to sampled time-points of the reference dataset (right).

Our newly identified erythropoietic MURK genes therefore perform key roles in red blood cell function, and their upregulation was validated in an independent model of red blood cell maturation.

### scRNA-Seq of mouse chimeras reveals the early cellular defects in Gata1 loss of function

The G1ER cell line represents an *in vitro* model, and the published differential gene expression data were from bulk microarray profiling, thus precluding any analysis of single-cell gene expression kinetics. We therefore turned to our recently reported Chimaera-Seq approach, whereby scRNA-Seq is coupled with mouse chimeric embryo technology, to define both cellular and molecular consequences of gene knock-outs *in vivo* (25, 31). We used our standard embryonic stem cells (ESCs) expressing a constitutive tdTomato (tdTom) fluorescent marker gene to generate a Gata1 knock-out line (see Methods). *Gata1*^-^ tdTom^+^ cells were injected into tdTom^-^ wild-type blastocyst and transferred into pseudo-pregnant females, resulting in chimeric embryos that we harvested at E8.5. Six chimeric embryos were pooled, dissociated into a single-cell suspension, and tdTom^+^ and tdTom^-^ cell fractions were sorted for scRNA sequencing. We obtained 8420 tdTom^-^ and 7944 tdTom^+^ cells passing quality control and assigned to a cell type, with an average of 4354 genes being detected per cell.

We then concatenated the chimera data with the Pijuan-Sala et al. (2019) reference dataset and mapped nearest neighbors (see Methods). We observed an overall homogeneous distribution of both mutant and wild-type fractions throughout the later time-points of the landscape, except for the erythroid branch. Indeed, we observed a block in the erythroid lineage of the mutant cells, which were over-represented in the start of the erythroid differentiation branch, while their wild-type counterparts were present throughout erythroid differentiation (Supplementary Figure 3). Identification of the nearest neighbours of chimeric cells within the reference dataset allowed their quick cell-type annotation, which we used to quantify the differences in the hemato-endothelial cell-type representation within the chimera fractions. This analysis confirmed a severe erythroid differentiation defect of the mutant cells (Figure 4C-E). When examining the reference dataset sampled-time point of the chimera nearest neighbours we also observed a temporal shift within the erythroid lineage, with tdTom^+^ mutant cells mapping to earlier time-points than their wildtype tdTom^-^ counterparts, further confirming a developmental block of the mutant cells (Figure 4D, E). In addition, we observed that this erythroid defect was coupled with an over-representation of cells with a megakaryocyte signature (Figure 4C).

The newly generated *Gata1*^-^ Chimaera-Seq data therefore not only recapitulated the expected block in erythroid maturation, but also revealed an expansion of the megakaryocytic lineage in the E8.5 yolk sac.

### The molecular program affected by Gata1 loss in early embryos

Although the role of Gata1 is well documented in developmental erythropoiesis (21, 23), the early molecular defects of Gata1 loss of function *in vivo* had not been reported. The Gata1 Chimaera-Seq dataset therefore presented an opportunity to dissect the early molecular program controlled by Gata1 *in vivo*. Having registered a defect in erythroid differentiation and an increase in the megakaryocytic lineage population, we performed differential gene expression testing between the chimera mutant and wild-type cells in these clusters (Supplementary Table 3).

Regarding the megakaryocytic subset, we observed upregulation of progenitor markers *Kit*, *Gata2* and *Myb* in the *Gata1*^-^ cells as well as lower expression of maturation genes for the megakaryocyte lineage *Gp5*, *Pf4*, *Mpl* and *Plek* (Figure 5A). Hyper-proliferative megakaryocyte progenitors, detected previously in *Gata1*^-^ E12.5 fetal livers, led to compromised platelet function, and were suggested to originate in the yolk sac (32). Our results showing over-production of megakaryocytic cells with impaired maturation characteristics in E8.5 *Gata1*^-^ chimera yolk sacs support this notion, and importantly place the megakaryocytic defect within the very early phase of megakaryocyte formation.

**Figure 5.**
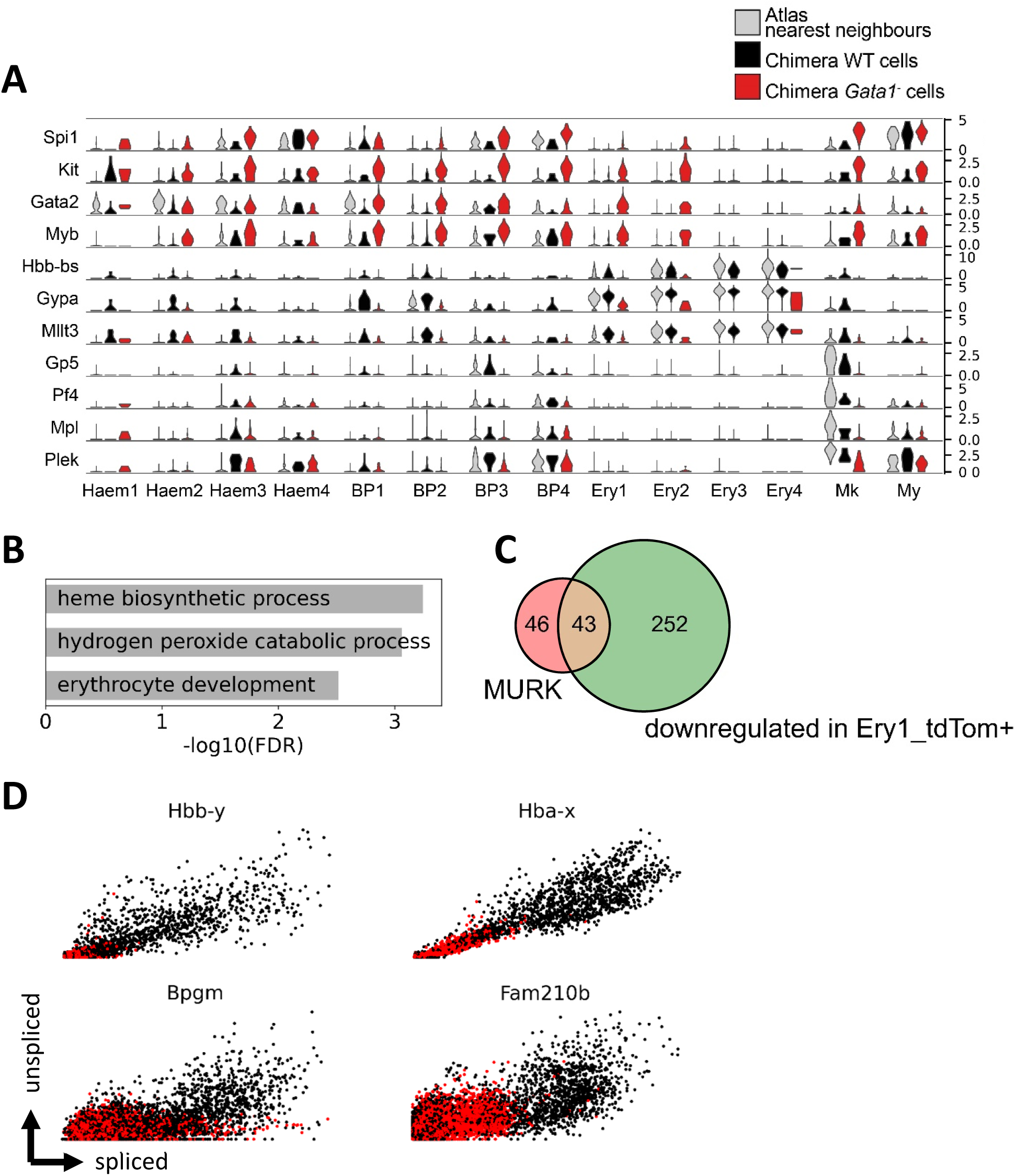
Gata1 chimaera assay reveals disruption of MURK genes and perturbed yolk sac hematopoiesis. A. Violin plots of representative genes differentially regulated in *Gata1*^-^ hematopoietic lineages. B. GO-term enrichment of genes downregulated in *Gata1*^-^ Ery1 cells compared to their WT counterparts in chimeras. C. Venn diagram showing overlap between MURK genes and genes downregulated in *Gata1*^-^ Ery1 cells D. Phase plots of MURK genes identified along erythroid differentiation, in E8.5 *Gata1*^-^ chimera datasets, colored by tdTom status.

Interestingly, all hemato-endothelial cell subsets displayed up-regulation of *Spi1* (coding for the PU.1 transcription factor) in the *Gata1*^-^ cell fraction compared to wild-type counterpart (FDR < 0.01; Figure 5A). Given the previously reported Gata1-PU.1 cross-repression in adult bone marrow (18) and in zebrafish embryonic hematopoiesis (33), we systematically assessed the effect of *Gata1* knockout in the mouse chimera lineages and observed that in *Gata1*^-^ cells, *Spi1* was specifically up-regulated in all hematopoietic sub-clusters, with a stronger effect on Mk and Ery1 subsets. (Supplementary Figure 3).

In the early erythroid subset, Ery1, we again noted that the mutant cells displayed increased expression of genes characteristic of a progenitor signature. Conversely, erythroid maturation hallmark genes such as *Hbb-bs* and *Gypa* were downregulated, along with the erythroid Gata1 target *Mllt3* (34; Figure 5A). GO-term enrichment analysis of genes downregulated in *Gata1*^-^ Ery1 cells revealed biological processes essential to red blood cell function (Figure 5B). Furthermore, we also observed that 48% of the MURK genes identified in Figure 3 overlapped with these genes that fail to up-regulate in *Gata1*^-^ erythroid cells (Figure 5C; p < 10^-24^).

In addition to the failure of inducing genes associated with erythroid maturation, single cell resolution molecular analysis also revealed a striking failure to downregulate genes associated with alternative lineage programs such as Pu.1, consistent with the notion that the earliest wave of primitive hematopoiesis produces erythroid cells, megakaryocytes and macrophages, with evidence for at least bipotential progenitor cells (35).

### The late erythroid increase in expression rate is downstream of Gata1 function

Having generated the Chimaera-Seq single cell data for both wildtype and Gata1 knock-out cells, we next used the ratio of spliced/unspliced reads to explore differences in expression kinetics between the wildtype and mutant cells. As can be seen in Figure 5D, the previously defined MURK genes failed to display the increased rate of expression characteristic for the later stages of erythropoiesis in the mutant cells. The examples shown include the embryonic globin gene *Hbb-y*, as well as the *Fam210b* gene, coding for a putative mitochondrial protein recently implicated in erythroid differentiation (36; Figure 5D). This result confirms that the erythroid boost in expression forms part of the transcriptional program downstream of Gata1 function, although it does not demonstrate a direct regulatory role for Gata1.

However, preliminary modelling analysis suggests that the change observed in MURK gene dynamics is due to altered transcription rates (see Supplementary Note), indicating a close association of the coordinated late erythroid increase in transcription rate with the molecular program downstream of Gata1.

### A coordinated increase of expression rate during human fetal liver erythropoiesis

Having identified a coordinated increase in transcription rate during mouse yolk sac erythropoiesis, we next wanted to ascertain whether the same phenomenon could also be seen in human cells. Moreover, we were keen to explore an scRNA-Seq dataset generated by a different laboratory, to exclude any potential technical bias caused by our own experimental protocols. We therefore turned to a recently published comprehensive dataset of human fetal liver erythropoiesis (37), and extracted the 49,388 cells annotated to the four clusters encompassing human fetal liver erythropoiesis. When calculating scVelo-based differentiation vectors as well as latent time using the full gene set (see methods), both were reversed (Figure 6A, left plots), consistent with the mouse yolk sac results. We therefore again ran our pipeline to discover genes with a potential increase in expression rate along the differentiation pathway. The resulting 97 genes again contained archetypal erythroid genes such as the haemoglobin genes (Figure 6B), with overall gene ontologies demonstrating a functional role in erythropoiesis (Figure 6C, see also Supplementary Table 4). We then recalculated both the scVelo differentiation vectors as well as latent time after removing the fetal liver MURK genes. This revealed scVelo vectors that were consistent with the expected developmental progression (see Figure 6A, right plots). This analysis therefore demonstrates that complex expression kinetics apply broadly to erythropoiesis, and their identification can be used to amend the RNA velocity framework to prevent erroneous predictions.

**Figure 6.**
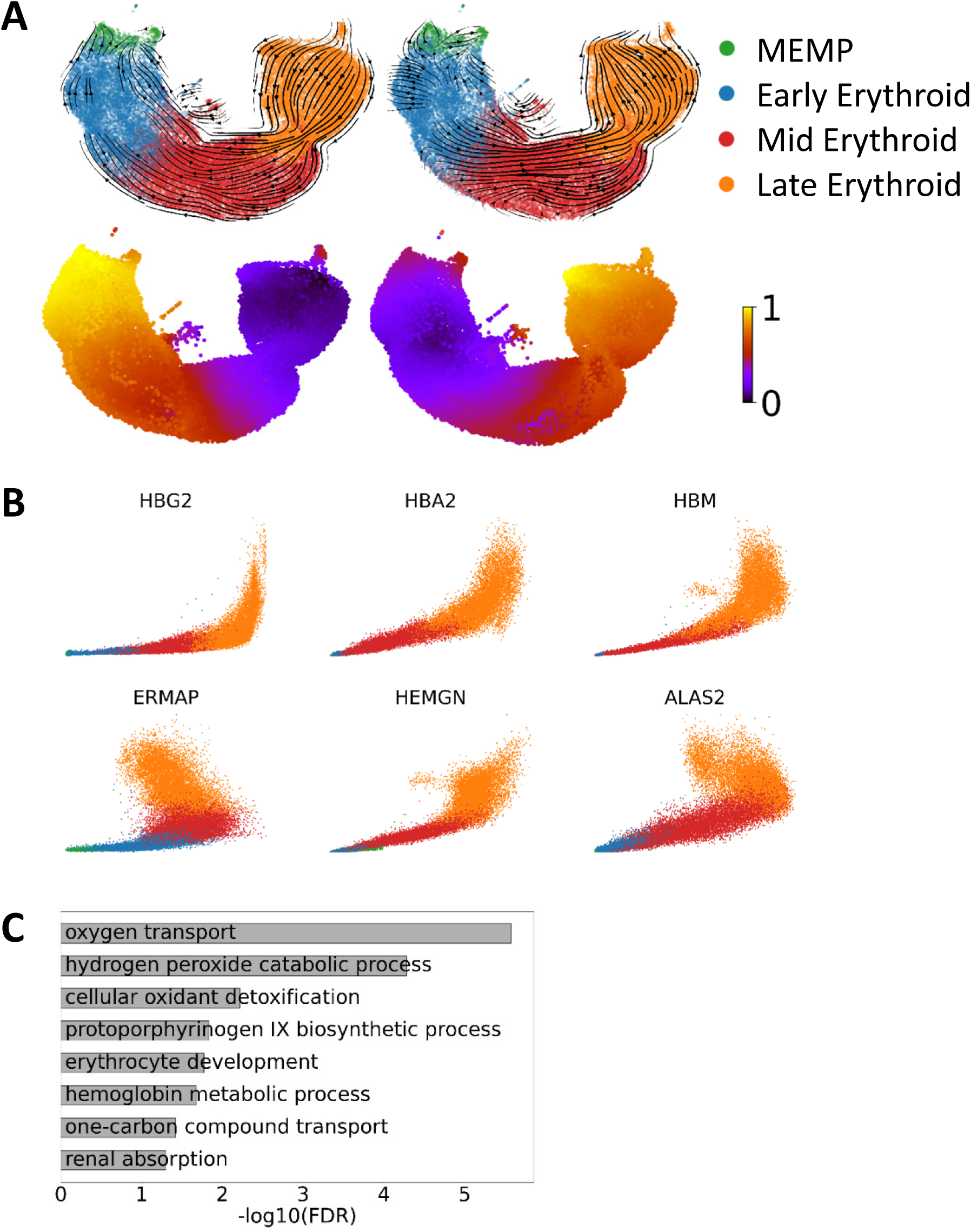
Concept of dual kinetics of gene expression is also revealed in human foetal liver hematopoiesis. A. UMAP representation of human fetal liver erythroid cell populations. The overlaying arrows result from applying the scVelo pipeline using all genes (left) or after MURK gene exclusion (right). Bottom UMAPs are colored by corresponding scVelo-inferred latent time. In order to facilitate comparison with the mouse data, a new clustering was performed on the erythroid cells, see Methods. MEMP: megakaryocyte-erythroid-mast cell progenitor. B. Phase plots of representative MURK genes identified in human fetal liver erythropoiesis single-cell RNAseq dataset. C. GO-term enrichment of MURK genes identified in human fetal liver erythropoiesis.

## Discussion

There is no doubt that single cell molecular profiling constitutes a transformative technology. It suffers however from the major drawback that cells need to be fixed in order to profile them, with the consequence that measurements are by necessity static snapshots. To decipher complex biological processes, however, temporal information is commonly required. The single cell RNA velocity concept raised the prospect of overcoming some of the limitations associated with static measurements, by providing a strategy that can infer future cellular states. The RNA velocity framework is based on an explicit model of transcriptional processes (transcription, splicing, degradation). The notion that physical parameters of gene expression can be deduced from single cell gene expression data had been explored before the single cell RNA velocity concept was introduced (38, 39). However, the scVelo implementation provided an attractive framework for estimating gene-specific expression parameters by taking advantage of the spliced versus unspliced read counts across large cell populations (14). Using erythropoiesis as an example, we show here that this current framework needs to be adapted to accommodate more complex expression kinetics. Importantly, our analysis revealed that sets of genes can show a coordinated increase in transcription rate along a differentiation pathway. Moreover, deletion of the key erythroid regulator Gata1 abrogated this coordinated change in expression dynamics, thus revealing this increase in transcription rate as an important feature of erythropoiesis. Of note, current RNA velocity frameworks consider only a single reason for the presence of introns, namely that a pre mRNA has not been fully processed. However, it is known that other processes such as intron retention can result in the presence of intronic sequences in otherwise fully processed cytoplasmic mRNA molecules (40, 41), thus suggesting that a more granular approach towards both the modelling and experimental analysis of spliced versus unspliced reads represents a promising avenue for future research.

Application of the single cell RNA velocity concept has commonly been “confirmatory”, whereby a differentiation path proposed by other means was shown to be consistent with RNA velocity inference. When we applied the RNA velocity framework to the entire mouse gastrulation atlas, some inferred vectors of differentiation agreed with our current understanding of developmental biology, but others disagreed. Deeper interrogation of predictions that conflicted with our current understanding of erythropoiesis showed that the RNA velocity predictions could not be correct, not only because they ran counter to the known expression changes that accompany red blood cell differentiation, but also because they contradicted the real-time sampling of the data. Our results thus highlight certain limitations of the current implementation of this framework for identification of novel trajectories. Importantly however, it is through our observation of the inconsistent predictions that we were led to identify the previously unrecognized dynamic nature of the transcriptional control of erythropoiesis. Moreover, it is plausible that coordinated increases in transcription rate midway through a differentiation process may operate more widely, as a powerful mechanism for stabilising a cell state. Our extension to the scVelo implementation reveals the presence of such time-dependent changes of gene expression parameters and retrieves the concerned MURK genes in developmental trajectories of interest.

As to the precise mechanisms, at this stage we can only confidently assert that this process occurs downstream of Gata1 during erythropoiesis. Of note, comprehensive analysis of the G1ER erythroid differentiation model has shown that Gata1-induced maturation triggers increased enhancer/promoter interactions for upregulated genes, and that the most highly enriched motif in the promoters of these genes are GATA sites (42). These observations are therefore consistent with the lineage-determining function of Gata1 involving a coordinated increase in expression kinetics of a set of genes important for red blood cell function.

Our observations regarding the Gata1 knock-out phenotype also warrant some discussion. With embryonically lethal phenotypes such as Gata1 knock-out, conventional analysis tends to be somewhat limited, since the embryos are dead because they have no red blood cells. By contrast, the Chimaera-Seq assay enables both quantification of cell numbers as well as characterisation of their molecular profiles. Moreover, there are no secondary effects caused by the dying embryo, because the wildtype host cells rescue overall fetal development, thus allowing a focussed analysis of cell-intrinsic molecular defects. One noteworthy observation from our data is that erythroid differentiation proceeds substantially beyond the stage where *Gata1* expression itself is first initiated, but fails to proceed to the late erythroid phase where expression of canonical red blood cell genes is greatly upregulated. However, gene expression prior to the differentiation block is not normal. In particular, we observed increased Spi1/Pu.1 in the Gata1 knock-out cells, consistent with the previously reported (18) but also disputed (43) antagonistic relationship between Gata1 and Pu.1.

Within haematopoiesis, Pu.1 is recognised as a key regulator of myeloid and T-cell lineages, but not erythroid cells, even though a role in the proliferation of immature erythroid progenitors has been reported (44, reviewed in 45). Upregulation of Pu.1 in our immature *Gata1* knock-out cells therefore suggests that these cells of the primitive haematopoietic lineage represent progenitors with multilineage potential, rather than being restricted to just the red cell lineage. Further evidence for this notion is provided by our observation that the reduction in erythroid cells in the *Gata1* knock-out is accompanied by an increase in megakaryocyte progenitors, consistent with a model whereby Gata1 levels influence the lineage choice decisions of a multipotent progenitor cell. Live cell tracking studies have suggested that the primary role of Gata1 and Pu.1 may be fate stabilization rather than fate choice (43). The increase in transcription rate of erythroid genes downstream of Gata1 would cohere with stabilizing the erythroid fate, thus suggesting that our results are consistent with roles in both fate choice and fate stabilization.

Our observation of an expanded pool of megakaryocyte progenitors may also be of direct relevance to our understanding of the pre-leukaemic transient myeloproliferative disease (TMD) that is prevalent in newborns with trisomy 21 (46). TMD is thought to arise when a fetal specific haematopoietic progenitor cell with trisomy 21 acquires a partial loss of function mutation in *GATA1*, resulting in a short form of GATA1 (GATA1s). TMD is characterized by expansion of immature megakaryocyte progenitors, and in 10 to 20% of cases transforms into malignant acute megakaryoblastic leukaemia (reviewed in 47). Over-expression of GATA1s in mouse models resulted in the identification of mid-gestation fetal liver megakaryocyte progenitors as uniquely sensitive to this mutant GATA1s form compared to their adult bone marrow counterparts (48). The over-represented population of immature megakaryocytic progenitors in our E8.5 *Gata1*^-^ chimeras may correspond to the developmental emergence of this transient precursor, TMD-initiating cell, in the yolk sac.

## Conclusions

Taken together, this study reports how the RNA velocity framework can be extended to delve into the transcriptional mechanisms of tissue differentiation, complemented with single cell resolution and *in vivo* analysis of Gata1 function, which revealed a number of previously unknown facets of this canonical regulator of red blood cell development.

## Methods

### scVelo implementation

#### Mouse atlas dataset

To obtain separated count matrices for spliced and unspliced mRNAs, we ran velocyto 0.17.17 (10) on the .bam files from the mouse atlas in Pijuan-Sala et al. 2019 (25; GEO accession number: GSE87038). We kept all cells that passed the QC as described in the original publication, but filtered out from downstream analysis the extraembryonic tissues: ExE endoderm, ExE ectoderm and Parietal endoderm as well as samples with no timepoint allocation (labelled as ‘mixed gastrulation’). To select highly variable genes (HVGs) we applied both the scanpy v1.5.1 and the scVelo v0.2.1 (14) pipelines. That is, we removed genes with less than 20 shared counts between spliced and unspliced counts, before normalising and log transforming the remaining genes. Then, we selected the top 2500 HVGs from each approach (resulting in a total of 4000, with 1000 overlapping genes) for further calculation of moments; while performing imputation using the top 30 nearest neighbours from the graph connectivities generated with the original UMAP coordinates from Pijuan-Sala et al. 2019. The velocity vectors were computed in dynamical mode rather than steady state.

#### Human dataset

We first downloaded raw reads from Popescu et al., 2019 (37; GEO accession number: GSE127980), and aligned them against the human genome hg19-3.0.0 with CellRanger v3.0.2 to generate the .bam files and obtain separated count matrices for spliced and unspliced mRNAs as described above. We filtered out cells with less than 3,550 counts, less than 900 genes and more than 6% mitochondrial counts. Again, we combined scapy and scVelo’s pipelines to select 1,500 HVGs to compute PCA coordinates and applied batch correction using the function reducedMNN from the batchelor package v1.4.0 (49), followed by the estimation of velocity vectors in the same way it was done for the mouse dataset.

### MOFA+ implementation

We ran MOFA+ v1.4.0 (26) using as input the two single cell experiment objects obtained from the spliced and unspliced counts independently. Each object was created in R using the scran v1.16.0 (50) library as follows: we started from the raw counts, normalized them with factor sizes obtained after pre-clustering, log transformed and reduced to 5000 HVG. We then switched to Python v3.7.4, where we regressed out the sample effect and scaled the object to generate a MOFA+ model with standard parameters. Finally, we used reducedMNN to correct the MOFA Factors for batch effects. The same objects used as MOFA input were used for PCA calculation in Figure 2A.

### MURK genes identification

To identify MURK genes, we considered the imputed counts resulting from the scVelo standard pipeline. Then, for each gene and each population among the Erythroid lineage, we calculated the unspliced versus spliced slope with a linear regression, as well as the standard error on the slope. In the mouse dataset we selected all genes for which the slope in Erythroid3 is significantly higher than the slope in Erythoid2 (according to a one-sided t-test p-value < 0.05), the average spliced counts in Erythroid3 is higher than the average spliced counts in every other population, and the slope in Erythroid3 positive. We found 89 genes that respect all these criteria.

In the human dataset, in order to obtain erythroid populations more comparable to our mouse data, we re-clustered the erythroid clusters (Figure 6A). We retained the population annotations from the original paper except for the Late Erythroid population, which we defined after performing Leiden clustering on the Umap coordinates. Specifically, we re-allocated a subset of the previously annotated Mid Erythroid population to Late Erythroid, in such a way that they have a similar numbers of cells. We then calculated the unspliced versus spliced slope with linear regression and identified MURK genes where the slope in Late Erythroid is significantly higher than the slope in Mid Erythroid. We found 97 genes respecting these criteria.

### Gene ontology enrichment analysis

We performed gene ontology enrichment analysis using the http://geneontology.org website comparing the MURK genes against all biological processes, with the default all Mus musculus genes in database as background set (51, 52). We ranked the processes by FDR.

### Overlap testing

Overlap was tested with Fisher exact test. We calculated the probability of having m = 55 genes of our n = 89 MURK genes mapping to the A = 1022 high response genes (out of N = 4195 genes) in the Wu et al., 2011 publication (GEO accession number: GSE30142) as the probability of randomly picking m elements of a specific type when randomly choosing n elements out of N, where the frequency of the special type is A/N.

### Gata1^-^ chimera dataset generation and analysis

#### Embryo collection

All procedures were performed in strict accordance to the UK Home Office regulations for animal research under the project license number PPL 70/8406. **Chimaera generation.** TdTomato-expressing mouse embryonic stem cells (ESC) were derived as previously described (25). Briefly, ESC lines were derived from E3.5 blastocysts obtained by crossing a male ROSA26tdTomato (Jax Labs – 007905) with a wildtype C57BL/6 female, expanded under the 2i+LIF conditions (53) and transiently transfected with a Cre-IRES-GFP plasmid (54) using Lipofectamine 3000 Transfection Reagent (ThermoFisher Scientific, #L3000008) according to manufacturer’s instructions. A tdTomato-positive, male, karyotypically normal line, competent for chimaera generation as assessed using morula aggregation assay, was selected for targeting *Gata1*. Two guides were designed using the http://crispr.mit.edu tool (guide 1: CGGCTACTCCACTGTGGCGG; guide 2: CGCTTCTTGGGCCGGATGAG) and were cloned into the pX458 plasmid (Addgene, #48138) as previously described (55). The obtained plasmids were then used to transfect the cells and single transfected clones were expanded and assessed for Cas9-induced mutations. Genomic DNA was isolated by incubating cell pellets in 0.1 mg/ml of Proteinase K (Sigma, #03115828001) in TE buffer at 50°C for 2 hours, followed by 5 min at 99°C. The sequence flanking the guide-targeted sites was amplified from the genomic DNA by polymerase chain reaction (PCR) in a Biometra T3000 Thermocycler (30 sec at 98°C; 30 cycles of 10 sec at 98°C, 20 sec at 58°C, 20 sec at 72°C; and elongation for 7 min at 72°C) using the Phusion High-Fidelity DNA Polymerase (NEB, #M0530S) according to the manufacturer’s instructions. Primers including Nextera overhangs were used (F-TCGTCGGCAGCGTCAGATGTGTATAAGAGACAGTCTACCCTGCCTCAACTGTG; R-GTCTCGTGGGCTCGGAGATGTGTATAAGAGACAGTCTTGTCTTGGGCAGGAACA), allowing library preparation with the Nextera XT Kit (Illumina, #15052163), and sequencing was performed using the Illumina MiSeq system according to manufacturer’s instructions. An ESC clone showing a 38 base-pair frameshift mutation in exon 4 resulting in the functional inactivation of *Gata1* were selected for injection into C57BL/6 E3.5 blastocysts. A total of 6 chimaeric embryos were harvested at E8.5, dissected, and single-cell suspensions were generated by TrypLE Express dissociation reagent (Thermo Fisher Scientific) incubation for 7-10 minutes at 37°C under agitation. Single-cell suspensions were sorted into tdTom+ and tdTom-samples using a BD Influx sorter with DAPI at 1µg/ml (Sigma) as a viability stain for subsequent 10X scRNA-seq library preparation (version 3 chemistry), and sequencing using an S1 flow cell in the Illumina Novaseq platform, which resulted in 8420 tdTom^-^ and 7944 tdTom^+^ cells that passed quality control (see “Single-cell RNA sequencing analysis” below).

#### Single-cell RNA sequencing analysis

Raw files were processed with Cell Ranger 3.0.2 using default mapping arguments. Reads were mapped to the mm10 genome and counted with GRCm38.92 annotation, including tdTomato sequence for chimera cells. Cell barcodes with expression profiles significantly different to the ambient mRNA expression profile were identified using emptyDrops (56), and cell barcodes with low complexity, i.e. low total mRNA counts and/or high mitochondrial proportion, were identified by fitting four-component bivariate mixture models to the log_10_-transformed total mRNA counts and percentage of mitochondrial counts, and selecting the components with high total mRNA and low mitochondrial percentage. Gene expression normalization and doublet cell barcodes were identified using the approach taken by Pijuan-Sala et al. (2019). Both spliced and unspliced count matrices were extracted using velocyto 0.17.17 (10).

#### Mapping to the reference dataset

We mapped the chimaera cells to the mouse atlas following almost exactly the procedure used in the original publication article to map the *Tal1* chimaera. First, we concatenated the mouse atlas and chimaera counts (both previously controlled for quality of the cells), normalized the resulting counts matrix with scran, computed HVGs and then applied multiBatchPCA, and reducedMNN with cosine normalization from batchelor (49) for batch effect correction within samples (where sample refers to a single lane of a 10x Chromium chip) as well as between datasets in order to extract a number of nearest neighbours between the mouse atlas and the chimaera using queryKNN from BiocNeighbors package v1.6.0.

#### Differential Gene Expression Analysis

For differential gene expression analysis, we took samples that included at least 7 cells per tdTom status per cell population (eg. Erythroid3). We ran the analysis in scanpy v1.5.1 (57) with Wilcoxon test and choosing 2 as fold change and 0.1 as false discovery rate thresholds.

## Funding

Research in the authors’ laboratories is supported by the Wellcome Trust, MRC, CRUK, Blood Cancer UK, NIH-NIDDK, the Sanger-EBI Single Cell Centre; by core support grants by the Wellcome Trust to the Cambridge Institute for Medical Research and Wellcome Trust-MRC Cambridge Stem Cell Institute; and by core funding from Cancer Research UK and the European Molecular Biology Laboratory. C.G. was funded by the Swedish Research Council (2017-06278), I.I. was funded by a British Heart Foundation studentship (FS/18/56/35177), S.G. was supported by a Royal Society Newton International Fellowship (NIF\R1\181950). This work was funded as part of a Wellcome Strategic Award (105031/D/14/Z) awarded to Wolf Reik, Berthold Göttgens, John Marioni, Jennifer Nichols, Ludovic Vallier, Shankar Srinivas, Benjamin Simons, Sarah Teichmann, and Thierry Voet.

## Authors’ contributions

M.B. performed scVelo implementations in mouse and human datasets, mathematical modelling, and analysis of Gata1 embryonic chimera dataset; I.I-R. assisted on the scVelo implementation in mouse datasets; I.I. performed Gata1 CRISPR/Cas9 targeting and expansion of the resulting mutant lines; S.G. performed quality controls of the Gata1 embryonic chimera dataset; S.G. and C.G. performed initial analysis of the Gata1 embryonic chimera dataset; C.G. designed and optimized the mutant chimera single-cell profiling experiments; B.G. wrote the initial draft of the manuscript; M.B., C.G., J.C.M. edited the manuscript; J.N., J.C.M., C.G. and B.G. supervised the study. All authors read and approved the final manuscript.

## Acknowledgements

We would like to thank Prof. Fabian Theis and Volker Bergen for discussions and valuable input on the scVelo implementation. We thank Prof. Ross Hardison for providing the list of Gata1-regulated genes from Wu et al. 2011 (Figure 4B). We thank William Mansfield and the Gurdon Institute animal facility for blastocyst injections, the Flow Cytometry Core Facility at the Cambridge Institute for Medical Research for cell sorting, Katarzyna Kania and the CRUK-CI genomics core for preparing the 10X libraries and for sequencing.

## Supplementary Figures

1. Dimensionality reduction with the first three principal components/MOFA factors using spliced reads alone (left), unspliced reads alone (middle) and both spliced and unspliced (right). Single-cell transcriptomes are colored by cell-type annotation; see Figure 1 for full legend.
2. Identification of MURK genes along yolk sac erythropoiesis. A. Phase plots of representative scVelo driver genes, with scVelo model prediction overlayed (see also Supplementary Table 1). B. Distribution of annotated cell type (top) and sampling time-point (bottom) along scVelo calculated latent time, using all genes (left panels) and after removing the MURK genes identified in Figure 3B-C.
3. Pijuan-Sala et al. (2019) layout highlighting nearest neighbours of *Gata1*^-^ chimeras. In red are nearest neighbours of tdTom+ mutant cells, in black those of tdTom- wildtype cells. To compare with Figure 1A.
4. Impact of Gata1 knockout on *Spi1*/PU.1 expression on the hematoendothelial cell types. X-axis: *Spi1* log_2_(fold-change) in *Gata1*^-^ vs WT chimera cells and Atlas nearest neighbours. Y-axis: log_10_(FDR).

## Supplementary Tables

1. Driver genes of the scVelo perdictions along erythroid differentiation, ranked by likelihood in the dynamic model.
2. List of mouse MURK genes identified in Figure 3B-C, ranked by calculated increase in slope value.
3. Differential Expression Analysis of Gata1^-^ tdTom^+^ vs WT tdTom^-^ chimera cells. For the Mk subset, given the low numbers of WT chimera cells present, the nearest neighbors from the reference Atlas dataset were included in the comparison. LFC: log fold change.
4. List of human MURK genes identified in Figure 6, ranked by calculated increase in slope value.

## Notes

### Competing Interest Statement

The authors have declared no competing interest.

## References

1. Akunuru S, Geiger H. Aging, Clonality, and Rejuvenation of Hematopoietic Stem Cells. Trends Mol Med. 2016;22(8):701–12.

2. Schultz MB, Sinclair DA. When stem cells grow old: phenotypes and mechanisms of stem cell aging. Development. 2016;143(1):3–14.

3. Mahadevaiah SK, Sangrithi MN, Hirota T, Turner JMA. A single-cell transcriptome atlas of marsupial embryogenesis and X inactivation. Nature. 2020;586(7830):612–7.

4. Gerber T, Murawala P, Knapp D, Masselink W, Schuez M, Hermann S, et al. Single-cell analysis uncovers convergence of cell identities during axolotl limb regeneration. Science. 2018;362(6413).

5. Wagner DE, Weinreb C, Collins ZM, Briggs JA, Megason SG, Klein AM. Single-cell mapping of gene expression landscapes and lineage in the zebrafish embryo. Science. 2018;360(6392):981–7.

6. Ton MN, Guibentif C, Gottgens B. Single cell genomics and developmental biology: moving beyond the generation of cell type catalogues. Curr Opin Genet Dev. 2020;64:66–71.

7. Borrett MJ, Innes BT, Jeong D, Tahmasian N, Storer MA, Bader GD, et al. Single-Cell Profiling Shows Murine Forebrain Neural Stem Cells Reacquire a Developmental State when Activated for Adult Neurogenesis. Cell Rep. 2020;32(6):108022.

8. Weinreb C, Rodriguez-Fraticelli A, Camargo FD, Klein AM. Lineage tracing on transcriptional landscapes links state to fate during differentiation. Science. 2020;367(6479).

9. Dahlin JS, Hamey FK, Pijuan-Sala B, Shepherd M, Lau WWY, Nestorowa S, et al. A single-cell hematopoietic landscape resolves 8 lineage trajectories and defects in Kit mutant mice. Blood. 2018;131(21):e1–e11.

10. La Manno G, Soldatov R, Zeisel A, Braun E, Hochgerner H, Petukhov V, et al. RNA velocity of single cells. Nature. 2018;560(7719):494–8.

11. Zhang Q, He Y, Luo N, Patel SJ, Han Y, Gao R, et al. Landscape and Dynamics of Single Immune Cells in Hepatocellular Carcinoma. Cell. 2019;179(4):829–45 e20.

12. Zhou W, Yui MA, Williams BA, Yun J, Wold BJ, Cai L, et al. Single-Cell Analysis Reveals Regulatory Gene Expression Dynamics Leading to Lineage Commitment in Early T Cell Development. Cell Syst. 2019;9(4):321–37 e9.

13. Kanton S, Boyle MJ, He Z, Santel M, Weigert A, Sanchis-Calleja F, et al. Organoid single-cell genomic atlas uncovers human-specific features of brain development. Nature. 2019;574(7778):418–22.

14. Bergen V, Lange M, Peidli S, Wolf FA, Theis FJ. Generalizing RNA velocity to transient cell states through dynamical modeling. Nat Biotechnol. 2020.

15. Grosveld F, van Assendelft GB, Greaves DR, Kollias G. Position-independent, high-level expression of the human beta-globin gene in transgenic mice. Cell. 1987;51(6):975–85.

16. Higgs DR, Wood WG, Jarman AP, Sharpe J, Lida J, Pretorius IM, et al. A major positive regulatory region located far upstream of the human alpha-globin gene locus. Genes Dev. 1990;4(9):1588–601.

17. Mettananda S, Gibbons RJ, Higgs DR. Understanding alpha-globin gene regulation and implications for the treatment of beta-thalassemia. Ann N Y Acad Sci. 2016;1368(1):16–24.

18. Zhang P, Behre G, Pan J, Iwama A, Wara-Aswapati N, Radomska HS, et al. Negative cross-talk between hematopoietic regulators: GATA proteins repress PU.1. Proc Natl Acad Sci U S A. 1999;96(15):8705–10.

19. McGrath K, Palis J. Ontogeny of erythropoiesis in the mammalian embryo. Curr Top Dev Biol. 2008;82:1–22.

20. Pevny L, Simon MC, Robertson E, Klein WH, Tsai SF, D’Agati V, et al. Erythroid differentiation in chimaeric mice blocked by a targeted mutation in the gene for transcription factor GATA-1. Nature. 1991;349(6306):257–60.

21. Pevny L, Lin CS, D’Agati V, Simon MC, Orkin SH, Costantini F. Development of hematopoietic cells lacking transcription factor GATA-1. Development. 1995;121(1):163–72.

22. Gutierrez L, Tsukamoto S, Suzuki M, Yamamoto-Mukai H, Yamamoto M, Philipsen S, et al. Ablation of Gata1 in adult mice results in aplastic crisis, revealing its essential role in steady-state and stress erythropoiesis. Blood. 2008;111(8):4375–85.

23. Fujiwara Y, Browne CP, Cunniff K, Goff SC, Orkin SH. Arrested development of embryonic red cell precursors in mouse embryos lacking transcription factor GATA-1. Proc Natl Acad Sci U S A. 1996;93(22):12355–8.

24. Shivdasani RA, Fujiwara Y, McDevitt MA, Orkin SH. A lineage-selective knockout establishes the critical role of transcription factor GATA-1 in megakaryocyte growth and platelet development. EMBO J. 1997;16(13):3965–73.

25. Pijuan-Sala B, Griffiths JA, Guibentif C, Hiscock TW, Jawaid W, Calero-Nieto FJ, et al. A single-cell molecular map of mouse gastrulation and early organogenesis. Nature. 2019;566(7745):490–5.

26. Argelaguet R, Arnol D, Bredikhin D, Deloro Y, Velten B, Marioni JC, et al. MOFA+: a statistical framework for comprehensive integration of multi-modal single-cell data. Genome Biol. 2020;21(1):111.

27. Storry JR, Joud M, Christophersen MK, Thuresson B, Akerstrom B, Sojka BN, et al. Homozygosity for a null allele of SMIM1 defines the Vel-negative blood group phenotype. Nat Genet. 2013;45(5):537–41.

28. Wu W, Cheng Y, Keller CA, Ernst J, Kumar SA, Mishra T, et al. Dynamics of the epigenetic landscape during erythroid differentiation after GATA1 restoration. Genome Res. 2011;21(10):1659–71.

29. Tsang AP, Visvader JE, Turner CA, Fujiwara Y, Yu C, Weiss MJ, et al. FOG, a multitype zinc finger protein, acts as a cofactor for transcription factor GATA-1 in erythroid and megakaryocytic differentiation. Cell. 1997;90(1):109–19.

30. Weiss MJ, Yu C, Orkin SH. Erythroid-cell-specific properties of transcription factor GATA-1 revealed by phenotypic rescue of a gene-targeted cell line. Mol Cell Biol. 1997;17(3):1642–51.

31. Guibentif C, Griffiths JA, Imaz-Rosshandler I, Ghazanfar S, Nichols J, Wilson V, et al. Diverse Routes toward Early Somites in the Mouse Embryo. Dev Cell. 2020.

32. Vyas P, Ault K, Jackson CW, Orkin SH, Shivdasani RA. Consequences of GATA-1 deficiency in megakaryocytes and platelets. Blood. 1999;93(9):2867–75.

33. Monteiro R, Pouget C, Patient R. The gata1/pu.1 lineage fate paradigm varies between blood populations and is modulated by tif1gamma. EMBO J. 2011;30(6):1093–103.

34. Pina C, May G, Soneji S, Hong D, Enver T. MLLT3 regulates early human erythroid and megakaryocytic cell fate. Cell Stem Cell. 2008;2(3):264–73.

35. Palis J. Hematopoietic stem cell-independent hematopoiesis: emergence of erythroid, megakaryocyte, and myeloid potential in the mammalian embryo. FEBS Lett. 2016;590(22):3965–74.

36. Kondo A, Fujiwara T, Okitsu Y, Fukuhara N, Onishi Y, Nakamura Y, et al. Identification of a novel putative mitochondrial protein FAM210B associated with erythroid differentiation. Int J Hematol. 2016;103(4):387–95.

37. Popescu DM, Botting RA, Stephenson E, Green K, Webb S, Jardine L, et al. Decoding human fetal liver haematopoiesis. Nature. 2019;574(7778):365–71.

38. Ezer D, Moignard V, Gottgens B, Adryan B. Determining Physical Mechanisms of Gene Expression Regulation from Single Cell Gene Expression Data. PLoS Comput Biol. 2016;12(8):e1005072.

39. Kim JK, Marioni JC. Inferring the kinetics of stochastic gene expression from single-cell RNA-sequencing data. Genome Biol. 2013;14(1):R7.

40. Edwards CR, Ritchie W, Wong JJ, Schmitz U, Middleton R, An X, et al. A dynamic intron retention program in the mammalian megakaryocyte and erythrocyte lineages. Blood. 2016;127(17):e24–e34.

41. Pimentel H, Parra M, Gee SL, Mohandas N, Pachter L, Conboy JG. A dynamic intron retention program enriched in RNA processing genes regulates gene expression during terminal erythropoiesis. Nucleic Acids Res. 2016;44(2):838–51.

42. Liu X, Chen Y, Zhang Y, Liu Y, Liu N, Botten GA, et al. Multiplexed capture of spatial configuration and temporal dynamics of locus-specific 3D chromatin by biotinylated dCas9. Genome Biol. 2020;21(1):59.

43. Hoppe PS, Schwarzfischer M, Loeffler D, Kokkaliaris KD, Hilsenbeck O, Moritz N, et al. Early myeloid lineage choice is not initiated by random PU.1 to GATA1 protein ratios. Nature. 2016;535(7611):299–302.

44. Choe KS, Ujhelly O, Wontakal SN, Skoultchi AI. PU.1 directly regulates cdk6 gene expression, linking the cell proliferation and differentiation programs in erythroid cells. J Biol Chem. 2010;285(5):3044–52.

45. Carotta S, Wu L, Nutt SL. Surprising new roles for PU.1 in the adaptive immune response. Immunol Rev. 2010;238(1):63–75.

46. Roberts I, Alford K, Hall G, Juban G, Richmond H, Norton A, et al. GATA1-mutant clones are frequent and often unsuspected in babies with Down syndrome: identification of a population at risk of leukemia. Blood. 2013;122(24):3908–17.

47. Bhatnagar N, Nizery L, Tunstall O, Vyas P, Roberts I. Transient Abnormal Myelopoiesis and AML in Down Syndrome: an Update. Curr Hematol Malig Rep. 2016;11(5):333–41.

48. Li Z, Godinho FJ, Klusmann JH, Garriga-Canut M, Yu C, Orkin SH. Developmental stage-selective effect of somatically mutated leukemogenic transcription factor GATA1. Nat Genet. 2005;37(6):613–9.

49. Haghverdi L, Lun ATL, Morgan MD, Marioni JC. Batch effects in single-cell RNA-sequencing data are corrected by matching mutual nearest neighbors. Nat Biotechnol. 2018;36(5):421–7.

50. Lun AT, McCarthy DJ, Marioni JC. A step-by-step workflow for low-level analysis of single-cell RNA-seq data with Bioconductor. F1000Res. 2016;5:2122.

51. Ashburner M, Ball CA, Blake JA, Botstein D, Butler H, Cherry JM, et al. Gene ontology: tool for the unification of biology. The Gene Ontology Consortium. Nat Genet. 2000;25(1):25–9.

52. The Gene Ontology C. The Gene Ontology Resource: 20 years and still GOing strong. Nucleic Acids Res. 2019;47(D1):D330–D8.

53. Ying QL, Wray J, Nichols J, Batlle-Morera L, Doble B, Woodgett J, et al. The ground state of embryonic stem cell self-renewal. Nature. 2008;453(7194):519–23.

54. Wray J, Kalkan T, Gomez-Lopez S, Eckardt D, Cook A, Kemler R, et al. Inhibition of glycogen synthase kinase-3 alleviates Tcf3 repression of the pluripotency network and increases embryonic stem cell resistance to differentiation. Nat Cell Biol. 2011;13(7):838–45.

55. Ran FA, Hsu PD, Wright J, Agarwala V, Scott DA, Zhang F. Genome engineering using the CRISPR-Cas9 system. Nat Protoc. 2013;8(11):2281–308.

56. Lun ATL, Riesenfeld S, Andrews T, Dao TP, Gomes T, participants in the 1st Human Cell Atlas J, et al. EmptyDrops: distinguishing cells from empty droplets in droplet-based single-cell RNA sequencing data. Genome Biol. 2019;20(1):63.

57. Wolf FA, Angerer P, Theis FJ. SCANPY: large-scale single-cell gene expression data analysis. Genome Biol. 2018;19(1):15.

