## Supplementary Figures for "Coordinated Changes in Gene Expression Kinetics Underlie both Mouse and Human Erythroid Maturation"

Supplementary Figure 1. Comparing variability-based dimensionality reduction methods

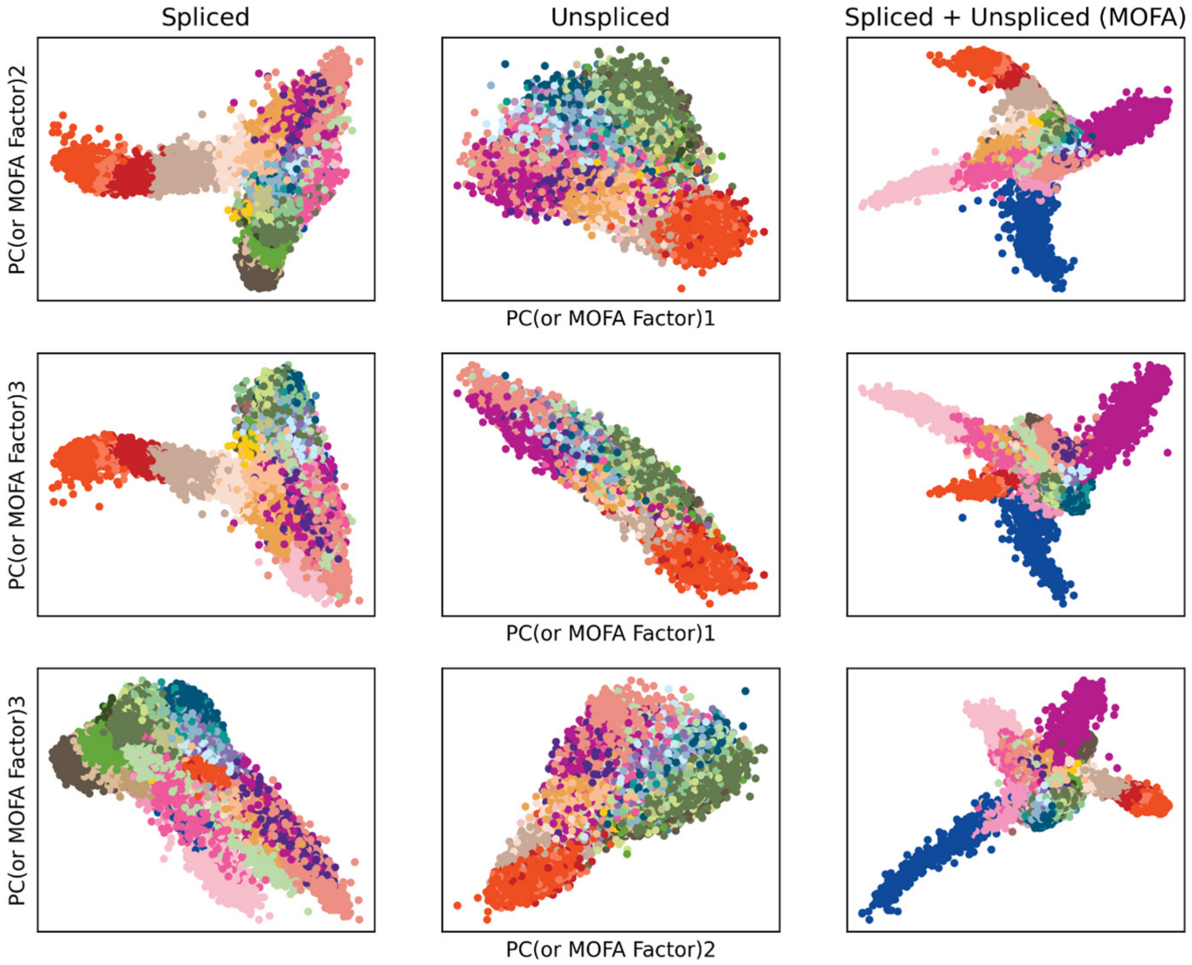

Supplementary Figure 2. Identification of MURK genes along yolk sac erythropoiesis

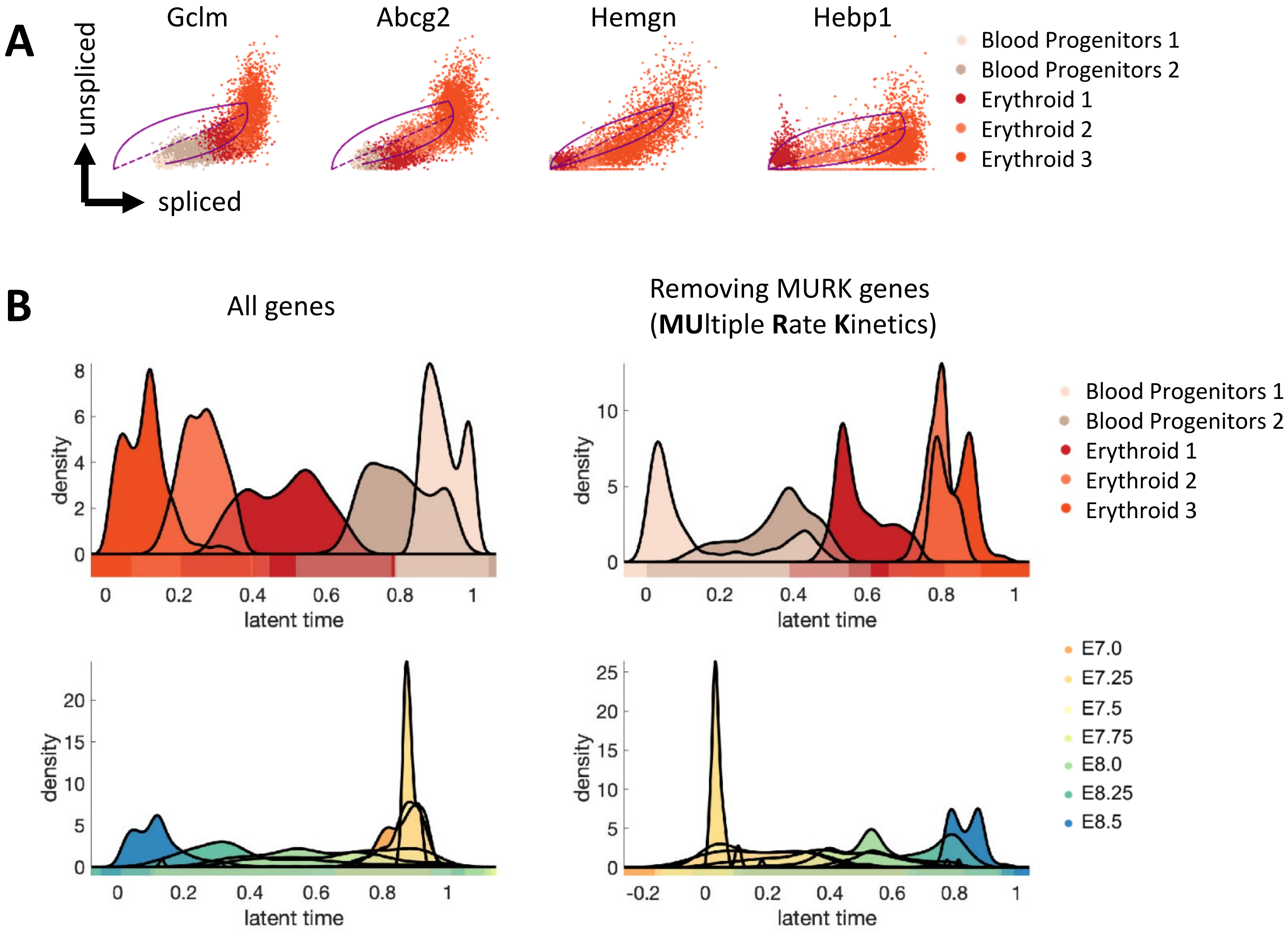

Supplementary Figure 3. Contribution to Chimera cells on the overall Atlas UMAP projection

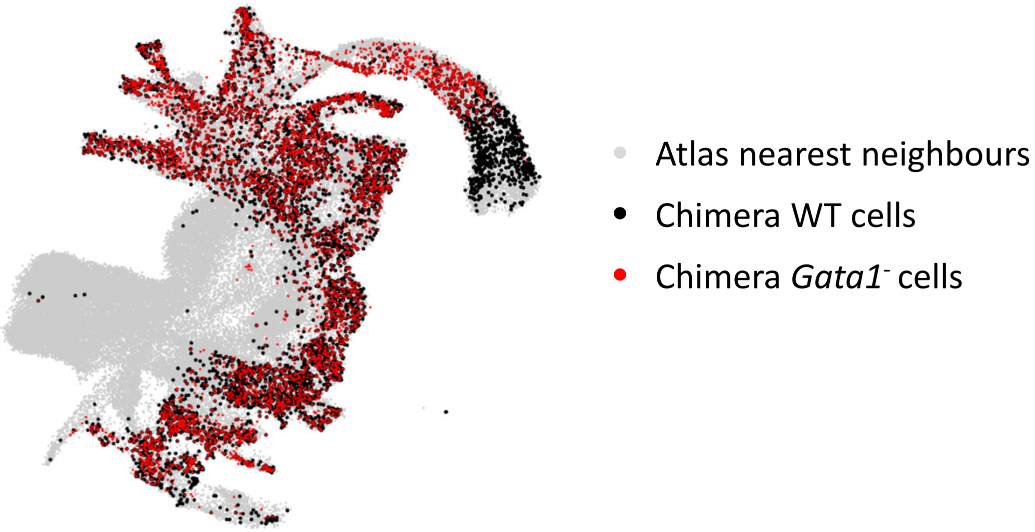

Supplementary Figure 4. Spi1 is upregulated in *Gata1*$^-$ yolk sac hematopoiesis

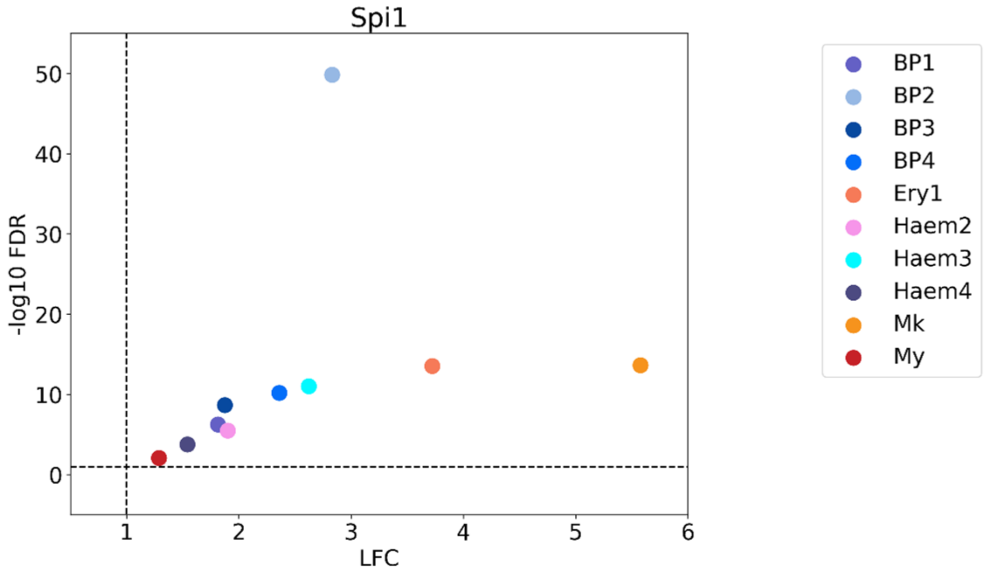
