## Supplementary Note for "Coordinated Changes in Gene Expression Kinetics Underlie both Mouse and Human Erythroid Maturation"

**Mathematical modelling**

We considered from the RNA velocity equations (La Manno et al. 2018, Bergen et al. 2020):

$$\frac{\mathrm{d}u(t)}{\mathrm{d}t} = \alpha(t) - \beta(t)u(t) \qquad (1)$$

$$\frac{\mathrm{d}s(t)}{\mathrm{d}t} = \beta(t)u(t) - \gamma(t)s(t) \qquad (2)$$

In these equations, *u(t)* and *s(t)* represent the number of unspliced and spliced counts over pseudotime respectively, while $\alpha(t)$, $\beta(t)$ and $\gamma(t)$ represent the three gene-specific kinetic parameters: rate of transcription, rate of splicing and rate of degradation. Given that these parameters are in principle also a function of pseudotime, we next asked the questions whether they are indeed pseudotime dependant or could be considered constant along the erythropoietic developmental trajectory. The parameters' functional dependence is unknown, but as argued in the text, they cannot be constant over the trajectory, otherwise the scVelo software would be able to fit the phase plot correctly. We thus assumed the easiest possible functional dependence that could represent a switch in kinetics, that is, the parameters are step functions with only one step, or equivalently:

$$\alpha_i(t) = \begin{cases} \alpha 1_i & t < \bar{t}_i \\ \alpha 2_i & t \geq \bar{t}_i \end{cases}$$

$$\beta_i(t) = \begin{cases} \beta 1_i & t < \bar{t}_i \\ \beta 2_i & t \geq \bar{t}_i \end{cases} \qquad (3)$$

$$\gamma_i(t) = \begin{cases} \gamma 1_i & t < \bar{t}_i \\ \gamma 2_i & t \geq \bar{t}_i \end{cases}$$

Where $\bar{t}_i$ is the specific latent time where the switch happens, considered the same for all parameters, and $\alpha 1_i$, $\alpha 2_i$, $\beta 1_i$, $\beta 2_i$, $\gamma 1_i$ and $\gamma 2_i$ are positive constants. Subscript *i* accounts for the different genes, treated as completely independently. All the parameters switch at the same pseudotime per gene. The constants and the switching point are parameters to be estimated. The switching time is allowed to vary between pseudotime 0.6-0.8 because, according to Figure 3E, at this point Erythroid3 emerge, and we visually see a switch in kinetics corresponding to the emergence of Erythroid3.

To calibrate equations (1) - (3) to our atlas dataset, we used as input the inputed unspliced and spliced counts, and we explicitly modelled the pseudotime as the latent time recovered after removing the MURK genes. Counts where binned in 10 bins according to the latent time and such that each bin contains the same number of cells. We thus took the average of each bin and used it as observable to model. More precisely, we built the cost function:

$$C_i(\alpha 1_i,\ \alpha 2_i,\ \beta 1_i,\ \beta 2_i,\ \gamma 1_i,\ \gamma 2_i,\ \delta_i,\ t_i) = \sum_{j=1}^{10}\left(\left(\frac{u_i(b_j)-u_i^o(b_j)}{\delta_i}\right)^2+\left(\frac{s_i(b_j)-s_i^o(b_j)}{\delta_i}\right)^2+2\log(\delta_i)\right) \quad (4)$$

Where $u_i(b_j)$ and $u_i^o(b_j)$ are the values of the modelled and the observed spliced counts at bin $b_j$, $u_i(b_j)$ and $u_i^o(b_j)$ are the values of the modelled and the observed spliced counts at bin $b_j$ and $\delta_i$ is the estimated standard error, same per bin, but estimated differently for each gene. The parameters' best values were calculated according to maximum likelihood estimation, that this upon minimising (4) via the *lsqnonlin* Matlab tool. Figure below shows an example of such fit to the top driver-gene, Smim1:

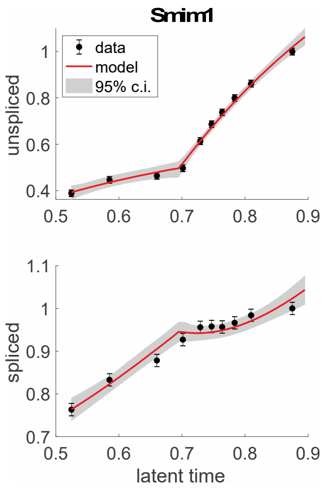

We next asked whether it is necessary to assume that all parameters switch values, when they switch at all. To address this question, we performed model selection, that is, for each gene independently, we fit these 8 models:

Model 1- all parameters are constant

Model 2- only transcription changes

Model 3- only splicing changes

Model 4- only degradation changes

Model 5- only transcription and splicing change

Model 6- only transcription and degradation change

Model 7- only splicing and degradation change

Model 8- all parameters change

We then ranked the models for each gene based on their performance, quantified by the corrected Akaike index A (Akaike 1978), which considers fit closeness to the *n*=20 datapoints (S, square sum of residuals), but penalises for the number of parameters used *k*. More precisely, for gene *i* and model *j*:

$$A_{i,j} = 2\,k_j + n\log\frac{S_{i,j}}{n} + 2\,\frac{k^2 + k}{n - k - 1}$$

We then ranked the indexes $A_{i,j}$ from lowest (model that contains more information) to the highest. Finally, for each model we calculated the average rank across genes, as in the following table:

| Model | Average Rank |
| --- | --- |
| Model 2 | 2.95 |
| Model 5 | 3.03 |
| Model 6 | 3.3 |
| Model 7 | 4.64 |
| Model 1 | 4.78 |
| Model 8 | 4.98 |
| Model 3 | 5.73 |
| Model 4 | 6.56 |

Table shows that the top model predicts a change in transcription over latent time, and in general the top 3 models are the models for which transcription at least changes. Worst model is Model 4, where only degradation changes.

Overall, our analysis suggests that at least the transcription rate is not constant over time.

**Reference**

Akaike, H. (1973), "Information theory and an extension of the maximum likelihood principle”
